# Computational analysis of memory consolidation following inhibitory avoidance (IA) training in adult and infant rats: critical roles of CaMKIIα and MeCP2

**DOI:** 10.1101/2022.03.30.486426

**Authors:** Yili Zhang, Paul Smolen, Cristina M. Alberini, Douglas A. Baxter, John H. Byrne

## Abstract

Key features of long-term memory (LTM), such as its stability and persistence, are acquired during processes collectively referred to as consolidation. The dynamics of biological changes during consolidation are complex. In adult rodents, consolidation exhibits distinct periods during which the engram is more or less resistant to disruption. Moreover, the ability to consolidate memories differs during developmental periods. Although the molecular mechanisms underlying consolidation are poorly understood, the initial stages rely on interacting signaling pathways that regulate gene expression, including brain-derived neurotrophic factor (BDNF) and Ca^2+^/calmodulin-dependent protein kinase II α (CaMKIIα) dependent feedback loops. We investigated the ways in which these pathways may contribute to developmental and dynamical features of consolidation. A computational model of molecular processes underlying consolidation following inhibitory avoidance (IA) training in rats was developed. Differential equations described the actions of CaMKIIα, multiple feedback loops regulating BDNF expression, and several transcription factors including methyl-CpG binding protein 2 (MeCP2), histone deacetylase 2 (HDAC2), and SIN3 transcription regulator family member A (Sin3a). This model provides novel explanations for the (apparent) rapid forgetting of infantile memory and the temporal progression of memory consolidation in adults. Simulations predict that dual effects of MeCP2 on the expression of *bdnf*, and interaction between MeCP2 and CaMKIIα, play critical roles in the rapid forgetting of infantile memory and the progress of memory resistance to disruptions. These insights suggest new potential targets of therapy for memory impairment.

**Author Summary:** Long-term memories (LTMs) are enduring and resistant to disruption These features are acquired *via* processes collectively referred to as consolidation. In adults, the initial stages of consolidation follow complex dynamics that are believed to emerge from interacting biochemical signaling pathways [1], including BDNF and CaMKIIα dependent feedback loops. Similarly, the acquisition of ability to consolidate memory in infantile animals is believed to emerge from the functional maturation of these molecular pathways [2]. Here, the ways in which these pathways contribute to consolidation were investigated using a computational model. This model provides novel explanations for the apparent rapid forgetting of infantile memory and for development of resistance to disruption during memory consolidation.

## Introduction

The transformation of an initially fragile memory into a stable long-term memory (LTM) is known as consolidation and can require days to complete [1]. Zhang et al. [3] developed a computational model of a hippocampal brain-derived neurotrophic factor (BDNF) - cAMP response element-binding protein (CREB)-CCAAT-enhancer-binding protein (C/EBPβ) positive feedback loop activated by inhibitory avoidance (IA) training in rats. Simulations predicted that the dynamics of the BDNF-CREB-C/EBPβ feedback loop have a significant effect on consolidation of long-term IA memory. Here, the Zhang et al. [3] model was extended to include additional signaling cascades:

1. Regulation of Ca^2+^/calmodulin-dependent protein kinase II alpha (CaMKIIα) is believed to be critical for the formation of IA memory. Based on empirical findings [4–6], a BDNF–CaMKIIα-BDNF feedback loop was included in the revised model.
2. The model adopted a ‘dual operation mode’ of regulation of BDNF by MeCP2 [7]. The effect of MeCP2 on *bdnf* expression has two components: activation by MeCP2 alone, and repression by a MeCP2 / HDAC2 / Sin3a complex.

We used the revised model to gain insights into apparent rapid decay of infantile memory. In contrast to memories formed in adults that will be stable and long-lasting, infantile memories formed early during development are usually more labile in that they commonly are no longer expressed days after learning, although the latent memory trace may be reinstated by some specific protocols or artificial reactivation [8–11]. In the hippocampus, many proteins related to synaptic plasticity have different basal expression levels in infants *vs*. adults. In infantile rats, the basal level of phosphorylated CaMKIIα (pCaMKIIα) is substantially lower than that of adult rats, whereas the basal level of phosphorylated CREB (pCREB) is substantially higher than that of adult [12]. Our revised model simulates IA memory formation with different basal levels of pCaMKIIα and pCREB.

We also used the model to study the development of resistance to disruption during memory consolidation following IA training. The consolidation of memory is a process of developing resistance to disruption [1]. Initially, memory is vulnerable to protein synthesis inhibitors (PSIs), but it becomes resistant over time, hence at the later stage of consolidation. IA memory at 7 days after learning can be impaired by PSI applied at 24 h after learning, but becomes resistant to PSI applied at 48 h after learning [5]. Empirical and computational studies suggest self-sustained positive feedback loops contribute to synaptic plasticity and memory formation and consolidation [3, 5, 13-24]. After positive feedback is activated by learning, its strength increases with time. Therefore, the later PSI is applied after learning, the more resistant is the positive feedback, and memory, to disruption [3].

However, resistance of memory to disruption does not always continuously grow with time post-learning. In Bekinschtein et al. [25], IA memory at day 7 was impaired by PSI added 12 h after training, but memory was resistant if PSI was added at earlier times. Some studies suggested that two distinct waves of protein synthesis during consolidation contribute to complex dynamics of resistance [25–27]. Zhang et al. [3] delayed the initiation of a BDNF feedback loop to generate two such waves, and consequent time windows of sensitivity to PSI. For this study, we simulated IA training and PSI effects in a more biologically realistic manner, and investigated biochemical processes that may underlie both the overall increased resistance to PSI with time, and the observed temporary decrease of resistance during a particular period after PSI application.

## Methods: Model development

The model of Zhang et al. [3] focused on the BDNF-CREB-C/EBPβ feedback loop in rat hippocampal neurons. In simulations, the feedback loop was initiated with a one-trial IA training-induced release of BDNF, which leads to rapid activation of the transcription factor CREB *via* phosphorylation, and a consequent increase in expression of the transcription factor C/EBPβ. Increased levels of C/EBPβ close the loop by increasing *bdnf* expression. We extended this model by adding several new components (Fig. 1A):

**Figure 1.**
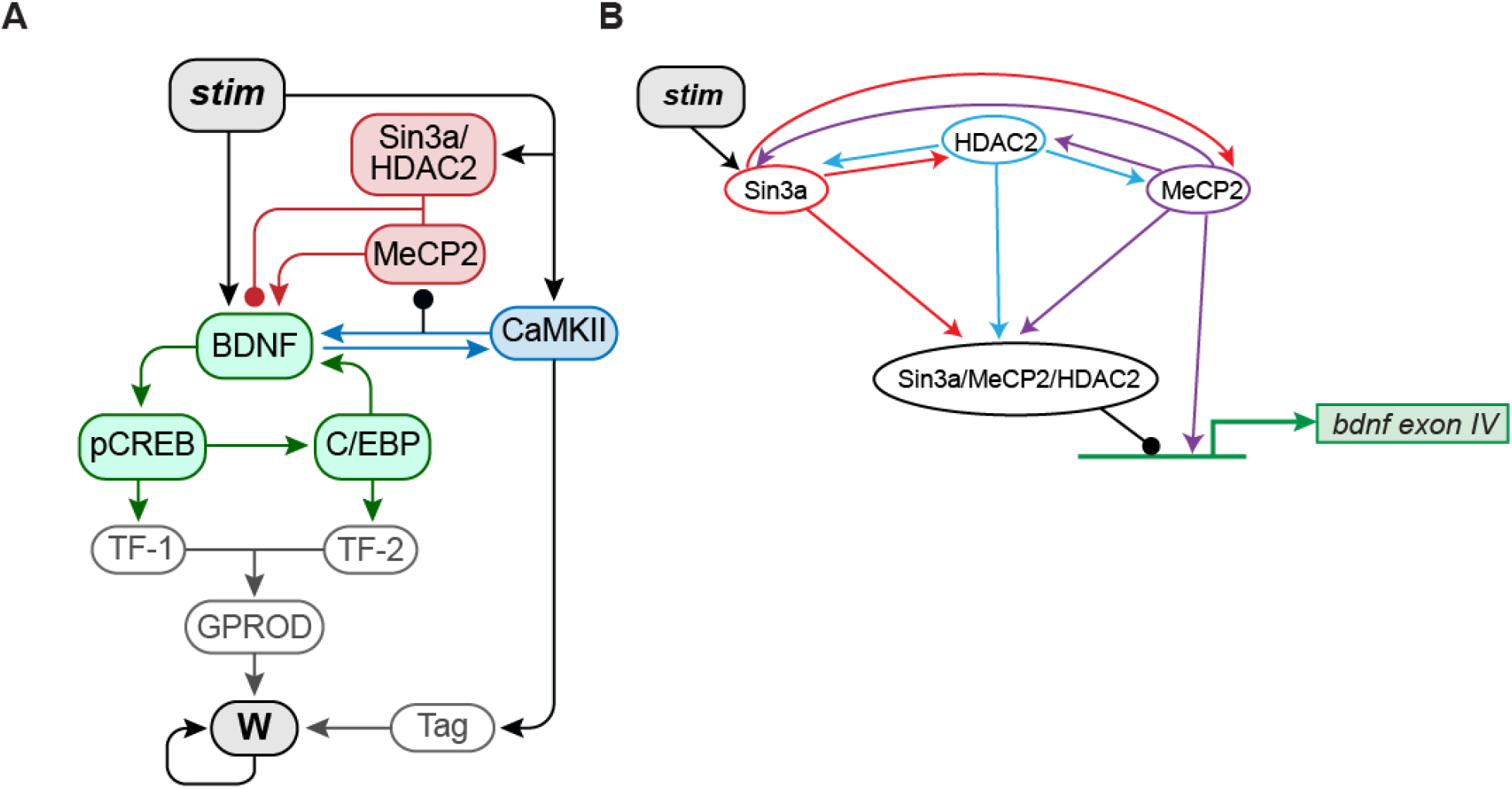
Schematic model of the signaling pathways essential to consolidation of IA long-term memory (LTM). (A) Signaling pathways with multiple feedback loops, including positive BDNF-CaMKIIα (blue) and BDNF-C/EBPβ (green) feedback loops that contribute to memory consolidation. MeCP2, HDAC2 and Sin3a bi-directionally regulate BDNF expression (red). Details and equations are given in the main text. (B) Scheme of interactions among bound Sin3a (red), MeCP2 (purple) and HDAC2 (blue), and their effects on the *bdnf* exon IV promoter. Arrowheads indicate activation, circular ends indicate repression.

1. <u>BDNF-CaMKIIα-BDNF positive feedback (Eqs. 1, 8-10)</u>. Empirical studies indicate that active CaMKIIα, i.e., phosphorylated CaMKIIα (pCaMKII) in the rat hippocampus remains increased for at least 20 h after IA training [4–5]. The increase of pCaMKIIα is significantly reduced by anti-BDNF treatment [5], which in turn blocks memory formation [4], suggesting activation of CaMKIIα is at least partially dependent on BDNF and accompanies memory formation. pCaMKIIα, in turn, supports extracellular release of BDNF and long-term changes in dendritic structure at CA3-CA1 hippocampal synapses [6]. Together, these results suggest autocrine BDNF-TrkB-CaMKIIα-BDNF signaling is important for synaptic plasticity and memory formation in the dorsal hippocampus. Thus, we included a BDNF-CaMKIIα-BDNF positive feedback loop (Fig. 1A). This loop does not require protein synthesis, thus the dynamics of this loop are faster compared to the BDNF-CREB-C/EBPβ feedback loop.
2. <u>Binding of MeCP2, Sin3a, and HDAC2 to the *bdnf* promoter (Eqs. 11-13)</u>. Modeling the dynamics of MeCP2, Sin3a, and HDAC2 binding to the *bdnf exon IV* promoter after IA training based on the empirical findings of Bambah-Mukku et al. [5] (Fig. 1). After IA training, distinct dynamics were observed for the binding of Sin3a, MeCP2 and HDAC2. Binding of Sin3a significantly increased within 30 min and decreased at 12 h, followed by a second increase at 48 h. In contrast, binding of MeCP2 and HDAC2 remained near the basal level for at least 12 h after IA training, but significantly increased at 48 h.
3. <u>Dual effects of MeCP2 on *bdnf* transcription (Eqs. 5-7)</u>. The action of MeCP2 on *bdnf* expression was modeled with two components: a constitutive function representing up-regulation of basal *bdnf* expression by free MeCP2, and a dynamic function to represent down-regulation by a MeCP2/HDAC2/Sin3a complex (Fig. 1). The effects of MeCP2 on gene expression are complex [28–29]. MeCP2 can act as a repressor or activator of BDNF transcription depending on context. In neurons, MeCP2 can act constitutively as a *bdnf* activator [30]. Constitutive MeCP2 overexpression enhances *bdnf* expression, and MeCP2 deletion inhibits *bdnf* expression [31]. Other studies suggest increased MeCP2 can act directly as a *bdnf* repressor [5, 32-33]. For example, termination of the BDNF positive feedback loop is associated with increased binding of MeCP2, HDAC2, and Sin3a to the *bdnf exon IV* promoter region at 48 h post IA training [5]. Li and Pozzo-Miller [7] proposed a “dual operation model” in which neuronal activity or other dynamical processes can switch the role of MeCP2 between activation and repression of *bdnf*. The association of MeCP2 with HDAC2-Sin3a could constitute such a switch component.
4. <u>Downstream signaling cascades to mediate synaptic plasticity (Eqs 14-21)</u>. These were based on the synaptic tagging and capture hypothesis of protein synthesis-dependent long-term potentiation [34].

### Equations for CaMKIIα-dependent regulation of the BDNF -CREB -C/EBPβ pathway

Eqs. 1-4 describe activation of the BDNF -CREB -C/EBPβ pathway after stimulation, modifed from those in Zhang et al. [3].

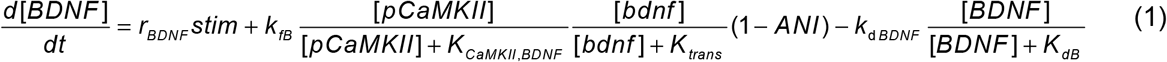

In Eq. 1, *stim* represents a stimulus that simulates IA training. *Stim* induces the initial activation of the BDNF pathway during training [35]. *Stim* is 0 before and after training, and increases to 30 μM/s for 1 min to simulate training. [*BDNF*] represents the concentration of BDNF released *via stim* or a CaMKIIα-dependent cascade. Released BDNF binds TrkB receptors to activate CREB and CaMKIIα. Active CaMKIIα in turn helps release more BDNF. *ANI* (anisomycin) remains at 0 in the absence of PSI and increases to 0.8 for 6 h to simulate PSI, based on empirical findings [5, 36].

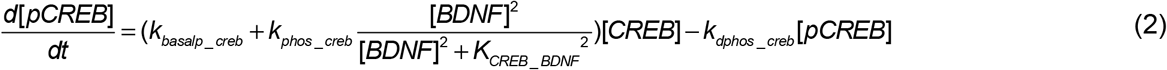

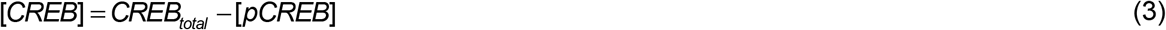

[*CREB_total_*] is the concentration of total CREB (Eq. 3). [*pCREB*] is phosphorylated CREB (pCREB) (Eq. 2). Empirical studies indicate that total CREB remains at its basal level for 20 h after IA training in rat hippocampus, whereas pCREB remains increased for more than 24 hours returning to baseline by 48 h after training [4–5]. The increase of pCREB is blocked by anti-BDNF treatment [5], suggesting the activation of CREB depends on BDNF.

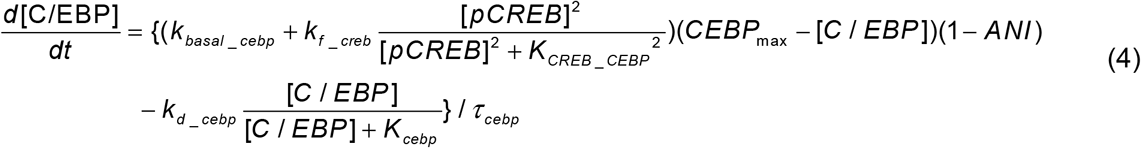

[*C/EBP*] is the total concentration of C/EBPβ protein (Eq. 4). Hill functions with coefficient of 2 are used to describe the effect of pCREB on *c/ebp* expression and the effect of C/EBPβ on *bdnf* expression, based on data suggesting these transcription factors commonly function as dimers. pCREB significantly increases 30 min after IA training [5], whereas *c/ebp*β mRNA and C/EBPβ protein do not increase until hours after training [5, 37]. The mechanism underlying the slow response of C/EBPβ is unclear. In the revised model, the arbitrary suppression of the effect of pCREB on C/EBPβ expression implemented in Zhang et al. [3] was removed. The response of C/BEPβ to pCREB is assumed slow. The parameter *τ_cebp_* in Eq. 4 determines the speed of response of C/EBPβ to pCREB. *τ_cebp_* was adjusted so that two waves of increase in the release of BDNF protein were induced after stimulus. The first was directly induced by stimulus, whereas the second was ~10 h after stimulus due to the activation of BDNF-C/EBPβ and BDNF-CaMKIIα feedback loops. *C/EBP_max_* is the maximum amount of C/EBPβ that can be produced within 48 h as described in Zhang et al. [3].

### Regulation of *bdnf* transcription by C/EBP, MeCP2, Sin3a, and HDAC2

Eqs. 5-7 describe the regulation of *bdnf* transcription by various factors. Eq. 5 is modified from the model of Zhang et al.[3]. Eqs. 6 and 7 are new equations.

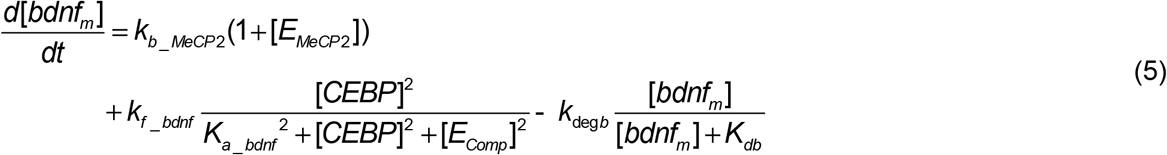

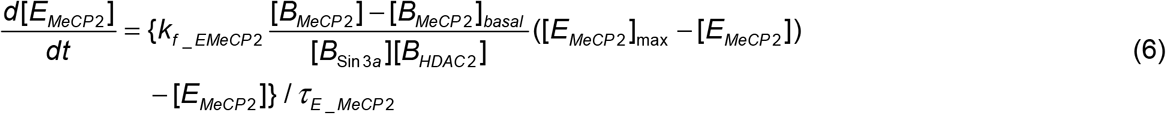

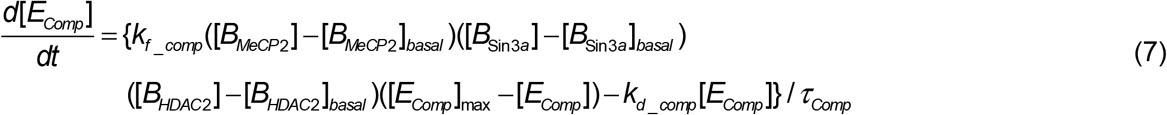

Transcription of *bdnf* is regulated by the activator C/EBPβ and three repressors: Sin3a, MeCP2, and HDAC2 [5]. In Eq. 5, the expression of *bdnf* mRNA is activated by [*C/EBP*], whereas *bdnf* is repressed by the effect of the Sin3a/MeCP2/HDAC2 complex, represented by the variable [*E_comp_*] (‘*E*’ representing ‘effect’; ‘*comp*’ representing ‘complex’). [*E_comp_*] increases due to increased binding of Sin3a ([*B_sin3a_*]), MeCP2 ([*B_MeCP2_*]), and HDAC2 ([*B_HDAC2_*]) (Eq. 7).

The effect of free MeCP2 on expression of *bdnf* is represented by [*E_MeCP2_*] (Eq. 5) (here ‘*MeCP2*’ represents free MeCP2 unbound to Sin3a or HDAC2). When the binding of MeCP2 to *bdnf exon IV* promoter is at the basal level, the basal translation rate is *k_b_MeCP2_* Overexpression or deletion of MeCP2 will increase or decrease [*E_MeCP2_*], respectively. [*E_MeCP2_*] increases with increased binding of MeCP2 alone ([*B_MeCP2_*] in Eq. 6) but decreases with increased binding of Sin3a ([*B_Sin3a_*]) and HDAC2 ([*B_HDAC2_*]), which bind MeCP2 to form the inhibitory Sin3a/MeCP2/HDAC2 complex (Eq. 6) (Fig. 1B). *τ_E_MeCP2_* and *τ_Comp_* are time constants governing how fast MeCP2 alone or MeCP2/Sin3a/HDAC2 complex respectively activate or repress the expression of *bdnf*.

### Equations for CaMKIIα regulation by BDNF

Eqs. 8-10 are new equations describing regulation of CaMKIIα. Empirical studies indicate that total CaMKIIα remains at its basal level after IA training in rat hippocampus, whereas pCaMKIIα is elevated for more than 24 h, returning to baseline by 48 h after training [4–5]. The increase of pCaMKIIα is significantly reduced by anti-BDNF treatment [5], suggesting this activation of CaMKIIα is dependent on BDNF (Eq. 8). However, pCaMKIIα is also increased earlier, 30 min after IA training, and this early increase is not reduced by anti-BDNF treatment [5]. Thus, in the model, *stim* in Eq. 8 is used to directly increase [*pCaMKII*] immediately after training. Also, the increase in pCaMKIIα at 12 h is not completely blocked by anti-BDNF treatment, which might be due to self-sustaining autophosphorylation of CaMKIIα [22, 38].

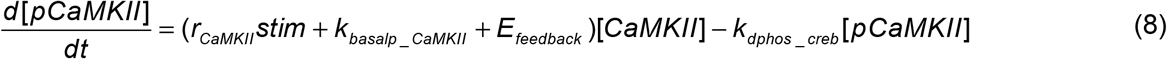

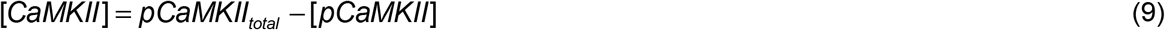

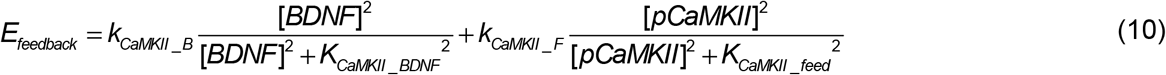

### Equations describing the dynamics of Sin3a, MeCP2 and HDAC2 binding

Eqs. 11-13 are new equations describing the binding of Sin3a, MeCP2 and HDAC2 to the *bdnf exon IV* promoter. Binding of MeCP2 does not significantly increase until 48 h after IA training [5], an effect that may be due to pCaMKIIα [39]. pCaMKIIα can phosphorylate MeCP2 on S421, decreasing binding of MeCP2 to methylated DNA [39]. The binding of Sin3a increases within 30 min after IA training [5]. The mechanisms underlying the changes in binding of Sin3a, MeCP2 and HDAC2 to the *bdnf* exon *IV* promoter are unclear. In the model, *stim* in Eq. 11 is used to directly increase [*B_sin3a_*] (the amount of bound Sin3a, Eq. 6) after training (Fig. 1B). Binding of Sin3a decreases at 12 h after IA training, followed by a second increase at 48 h, along with binding of MeCP2 and HDAC2. We hypothesize these similarities in binding dynamics reflect concurrent binding of Sin3a, MeCP2, and HDAC2 to form the Sin3a/MeCP2/HDAC2 complex. Thus increasing the binding of any two components (e.g., [*B_MeCP2_*] and [*B_sin3a_*]) will increase binding of the third (i.e., [*B_HDAC2_*]) (Eqs. 12, 13) (Fig. 1B). [*B_Sin3a_*]_max_, [*B_MeCP2_*]_max_ and [*B_HDAC2_*]_max_ represent the maximal binding of these components.

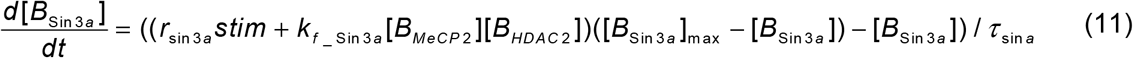

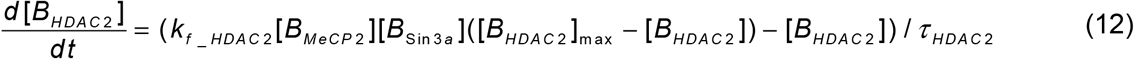

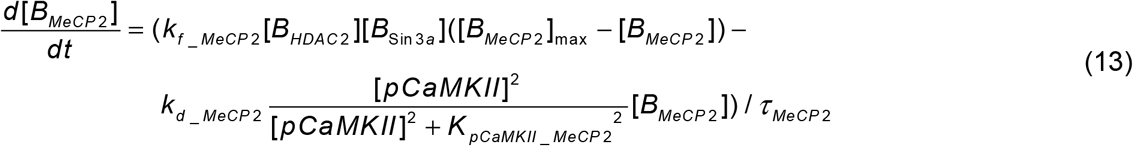

### Equations describing the synaptic tagging and capture system

Eqs. 14-21 are new equations describing the downstream synaptic tagging and capture system. These equations are adapted from [34, 40]. CaMKIIα activates a synaptic tag (Tag) in Eq. 14. Tag is required for synaptic ‘capture’ of protein synthesized from a generic gene (GPROD in Eq. 17) necessary for synaptic strengthening and LTM. Phosphorylated CREB activates a generic transcription factor TF-1 (Eq. 15), whereas C/EBPβ activates a second transcription factor TF-2 (Eq. 16) (Fig. 1A). The rate of synthesis of GPROD, assumed necessary for synaptic strengthening, increases with TF-1 and TF-2 activation (Eq. 17) (Fig. 1A). The rate of increase of a synaptic weight W (Eqs. 18-20) is determined by the product of Tag and GPROD (Fig. 1A). The model simulates long-term synaptic potentiation (LTP) (increases in W, corresponding to formation of LTM) but does not currently simulate long-term synaptic depression (LTD). Thus, regulation of W is modeled by increasing W when the product of GPROD and Tag increases above its basal level. A Heaviside function, denoted as ()^+^, is used to represent regulation by Tag and GPROD (Eq. 21). When the product of Tag and GPROD is below a basal level, [*Tag*]*_basal_ x* [*GPROD*]*_basal_*, Tag and GPROD do not affect W.

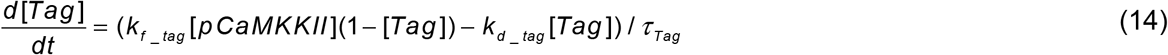

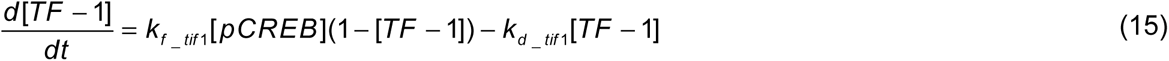

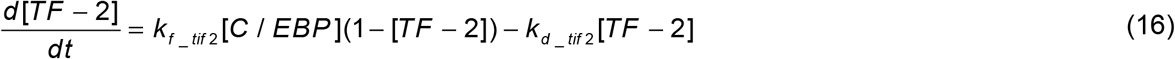

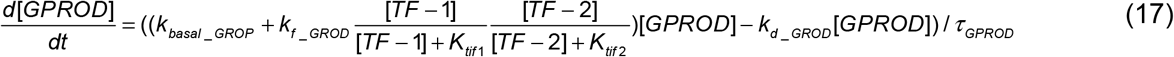

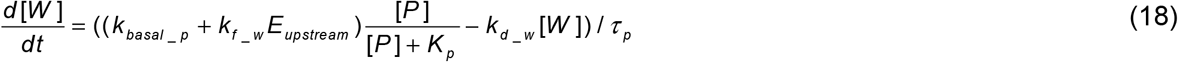

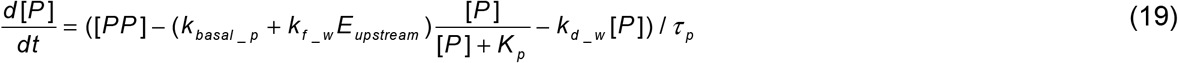

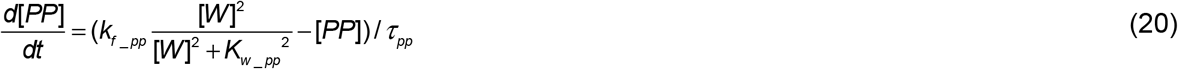

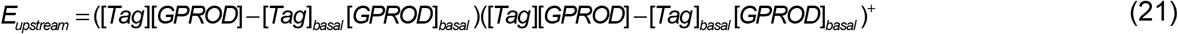

so that

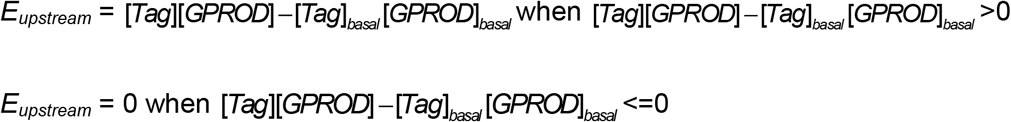

As in Smolen et al. [34, 40], the increase in W is also assumed to be limited by the availability of a precursor molecule P (Eq. 19). Moreover, in the current model, as a reactivating mechanism to maintain persistent memory (> 7 days), a simple positive feedback loop in which increased W tends to favor further growth, or stabilization, of W [22] is implemented. Increased W is assumed to feed back to increase the availability of P, through an intermediate pathway represented by PP (Eqs. 20, 19). This feedback can allow W to remain at 200% or higher of its unstimulated (basal) value, a threshold set in Zhang et al. [3]. Long-term memory is assumed to be consolidated if W remains at least 200% or higher of its basal value for more than 2 days.

The parameter values in Table 1 were adjusted so that the model replicated the characteristic features of dynamics of BDNF, *bdnf*, pCREB, C/EBPβ and pCaMKIIα, and the binding of Sin3a/MeCP2/HDAC2 after IA training, as described in Bambah-Mukku et al. [5], and also maintained W higher than 200% control to simulate IA memory consolidation (Table 1).

**Table 1.** Standard parameter values.

| Parameter name | Value |
| --- | --- |
| $r_{BDNF}$ | $0.3 \text{ s}^{-1}$ |
| $k_{fB}$ | $1.5 \times 10^{-3} \text{ mM/s}$ |
| $K_{CaMKII, BDNF}$ | $7.3 \times 10^{-3} \text{ mM}$ |
| $K_{trans}$ | $3 \times 10^{-2} \text{ mM}$ |
| $K_{dB}$ | $0.6 \text{ mM}$ |
| $k_{dBDNF}$ | $6.6 \times 10^{-4} \text{ mM/s}$ |
| $k_{basalp\_creb}$ | $3.2 \times 10^{-6} \text{ s}^{-1}$ |
| $k_{dphos\_creb}$ | $9.6 \times 10^{-5} \text{ s}^{-1}$ |
| $k_{phos\_creb}$ | $4 \times 10^{-3} \text{ s}^{-1}$ |
| $K_{CREB\_BDNF}$ | $2 \text{ mM}$ |
| $CREB_{total}$ | $0.085 \text{ mM}$ |
| $k_{basal\_cebp}$ | $1.6 \times 10^{-2} \text{ s}^{-1}$ |
| $k_{f\_cebp}$ | $2.2 \text{ s}^{-1}$ |
| $k_{d\_cebp}$ | $5.7 \times 10^{-3} \text{ mM/s}$ |
| $K_{CREB\_CEBP}$ | $0.19 \text{ mM}$ |
| $CEBP_{max}$ | $2.3 \times 10^{-1} \text{ mM}$ |
| $K_{cebp}$ | $7.6 \times 10^{-2} \text{ mM}$ |
| $\tau_{cebp}$ | $3000 \text{ s}$ |
| $k_{b\_MeCP2}$ | $1.6 \times 10^{-7} \text{ mM/s}$ |
| $k_{f\_bdnf}$ | $1.6 \times 10^{-4} \text{ mM/s}$ |
| $K_{a\_bdnf}$ | $3.2 \text{ mM}$ |
| $k_{degb}$ | $4.5 \times 10^{-5} \text{ mM/s}$ |
| $K_{db}$ | $0.6 \text{ mM}$ |
| $k_{f\_EMeCP2}$ | $0.34 \text{ mM}$ |
| $[E_{MeCP2}]_{max}$ | $5 \text{ mM}$ |
| $\tau_{E\_MeCP2}$ | $1.44 \times 10^4 \text{ s}$ |
| $k_{f\_comp}$ | $4.5 \text{ mM}$ |
| $k_{f\_comp}$ | $6.7 \times 10^{-3} \text{ mM}$ |
| $[B_{Sin3a}]_{basal}$ | $1 \text{ mM}$ |
| $[B_{HDAC2}]_{basal}$ | $1 \text{ mM}$ |
| $[I_{Comp}]_{max}$ | $5 \text{ mM}$ |
| $\tau_{Comp}$ | $3.6 \times 10^4 \text{ s}$ |
| $r_{CaMKII}$ | $0.13 \text{ s}^{-1}$ |
| $k_{basalp\_CaMKII}$ | $2.8 \times 10^{-6} \text{ mM/s}$ |
| $k_{\text{dphos\_CaMKII}}$ | $8 \times 10^{-5} \text{ s}^{-1}$ |
| $p\text{CaMKII}_{\text{total}}$ | 0.085 mM |
| $k_{\text{CaMKII\_B}}$ | $4 \times 10^{-5} \text{ mM/s}$ |
| $K_{\text{CaMKII\_BDNF}}$ | 0.28 mM |
| $k_{\text{CaMKII\_F}}$ | $3.1 \times 10^{-5} \text{ mM/s}$ |
| $K_{\text{CaMKII\_feed}}$ | 0.03 mM |
| $r_{\text{sin3a}}$ | $120000 \text{ mM}^{-1} \text{ s}^{-1}$ |
| $k_{\text{f\_Sin3a}}$ | 0.25 mM |
| $\tau_{\text{sin3a}}$ | 50000 s |
| $[B_{\text{Sin3a}}]_{\text{max}}$ | 5 mM |
| $k_{\text{f\_HDAC2}}$ | 0.25 mM |
| $[B_{\text{HDAC2}}]_{\text{max}}$ | 5 mM |
| $\tau_{\text{HDAC2}}$ | 125000 s |
| $k_{\text{f\_MeCP2}}$ | 0.25 mM |
| $[B_{\text{MeCP2}}]_{\text{max}}$ | 5 mM |
| $k_{\text{d\_MeCP2}}$ | 4 mM |
| $K_{\text{pCaMKII\_MeCP2}}$ | 0.011 mM |
| $\tau_{\text{MeCP2}}$ | $1 \times 10^4 \text{ s}$ |
| $k_{\text{f\_tag}}$ | $0.5 \text{ mM}^{-1} \text{ s}^{-1}$ |
| $k_{\text{d\_tag}}$ | $0.03 \text{ s}^{-1}$ |
| $\tau_{\text{Tag}}$ | 5 s |
| $k_{\text{f\_tif1}}$ | $1.2 \times 10^{-4} \text{ mM}^{-1} \text{ s}^{-1}$ |
| $k_{\text{d\_tif1}}$ | $3 \times 10^{-5} \text{ s}^{-1}$ |
| $k_{\text{f\_tif2}}$ | $2.4 \times 10^{-5} \text{ mM}^{-1} \text{ s}^{-1}$ |
| $k_{\text{d\_tif2}}$ | $6 \times 10^{-5} \text{ s}^{-1}$ |
| $k_{\text{basal\_GROP}}$ | $4 \times 10^{-4} \text{ mM/s}$ |
| $k_{\text{f\_GROD}}$ | 7.5 mM/s |
| $k_{\text{d\_GROD}}$ | $0.05 \text{ s}^{-1}$ |
| $K_{\text{tif1}}$ | 1 mM |
| $\tau_{\text{GPROD}}$ | 2000 s |
| $K_{\text{tif2}}$ | 1 mM |
| $k_{\text{basal\_p}}$ | 0.32 mM |
| $k_{f\_w}$ | 1680 mM |
| $K_p$ | 15.2 mM |
| $k_{d\_w}$ | $0.08 \text{ s}^{-1}$ |
| $[tag]_{basal}$ | 0.1 mM |
| $[GPROD]_{basal}$ | 0.12 mM |
| $K_{w\_pp}$ | 0.9 mM |
| $\tau_p$ | $2.5 \times 10^4 \text{ s}$ |
| $\tau_{pp}$ | $7.5 \times 10^5 \text{ s}$ |
| $k_{f\_pp}$ | 0.8 mM |

### Numerical methods

Fourth-order Runge-Kutta integration was used for integration of all differential equations with a time step of 3 s. Further time step reduction did not lead to significant improvement in accuracy. The steady-state levels of variables were determined after at least two simulated days, prior to any manipulations. The model was programmed in XPPAUT (http://www.math.pitt.edu/~bard/xpp/xpp.html) [41]. Source codes will be submitted to the ModelDB database [42] and to GitHub (https://github.com).

## Results

### Simulated variations of W following IA training with altered CaMKIIα and CREB basal activities

Memories formed early during development are usually more labile in that they commonly are no longer expressed days after learning, although the latent memory trace may be reinstated by some specific protocols [8-9, 12]. This lability is thought to be linked to infantile amnesia, the inability of adults to remember infantile experiences. Although multiple hypotheses have been proposed [8-10, 12, 43-44], the mechanisms of rapid forgetting of infantile memory and reinstatement of latent memory traces remain unclear. Travaglia et al. [12] point out some of the substantial differences in biological systems of developing and adult brains. In the dorsal hippocampus of infantile rats, the basal level of pCaMKIIα is substantially lower than that of adult rats, whereas the basal level of pCREB is substantially higher than in adult [12]. To simulate the effects of these changes on the consolidation of IA memory, we gradually increased the basal phosphorylation rate of CREB, *k_basalp_creb_* in Eq. 2, from 100% of its standard value in Table 1 to ~300%, by steps of 20%. At the same time, we gradually decreased the basal phosphorylation rate of CaMKIIα, *k_basalp_CaMKII_* in Eq. 8, from 100% of its standard value in Table 1 to 50%, by steps of 10% (arrows in Fig. 2A1). In all cases we first simulated several days of stimulus absence until the model reached a new equilibrium, and then simulated how these parameter changes affect the response to a stimulus. Basal binding of MeCP2, Sin3a and HDAC2 to the *bdnf* exon *IV* promoter was increased compared to control, due to the reduced basal pCaMKIIα, releasing its inhibition of MeCP2 binding. The increase in MeCP2 subsequently increased the binding of Sin3a and HDAC2. Figure 2A2 summarizes the results of these simulations with a 3-D plot of synaptic weight W as a function of both parameters *k_basalp_CaMKII_* and *k_basalp_creb_*. The light blue area represents W lower than 200% of basal level. Starting from the control simulation using the standard values of *k_basalp_creb_* and *k_basalp_CaMKII_* in Table 1, if the basal phosphorylation rate of CaMKIIα was reduced by more than 40%, *W* at day 7 always fell below the 200% threshold that represents consolidated LTM, even if *k_basalp_creb_* was substantially increased by more than 100% (light blue area, Fig. 2A2). These simulations suggest that decreased pCaMKIIα in infant animals plays a role in the rapid forgetting of infantile memory.

**Figure 2.**
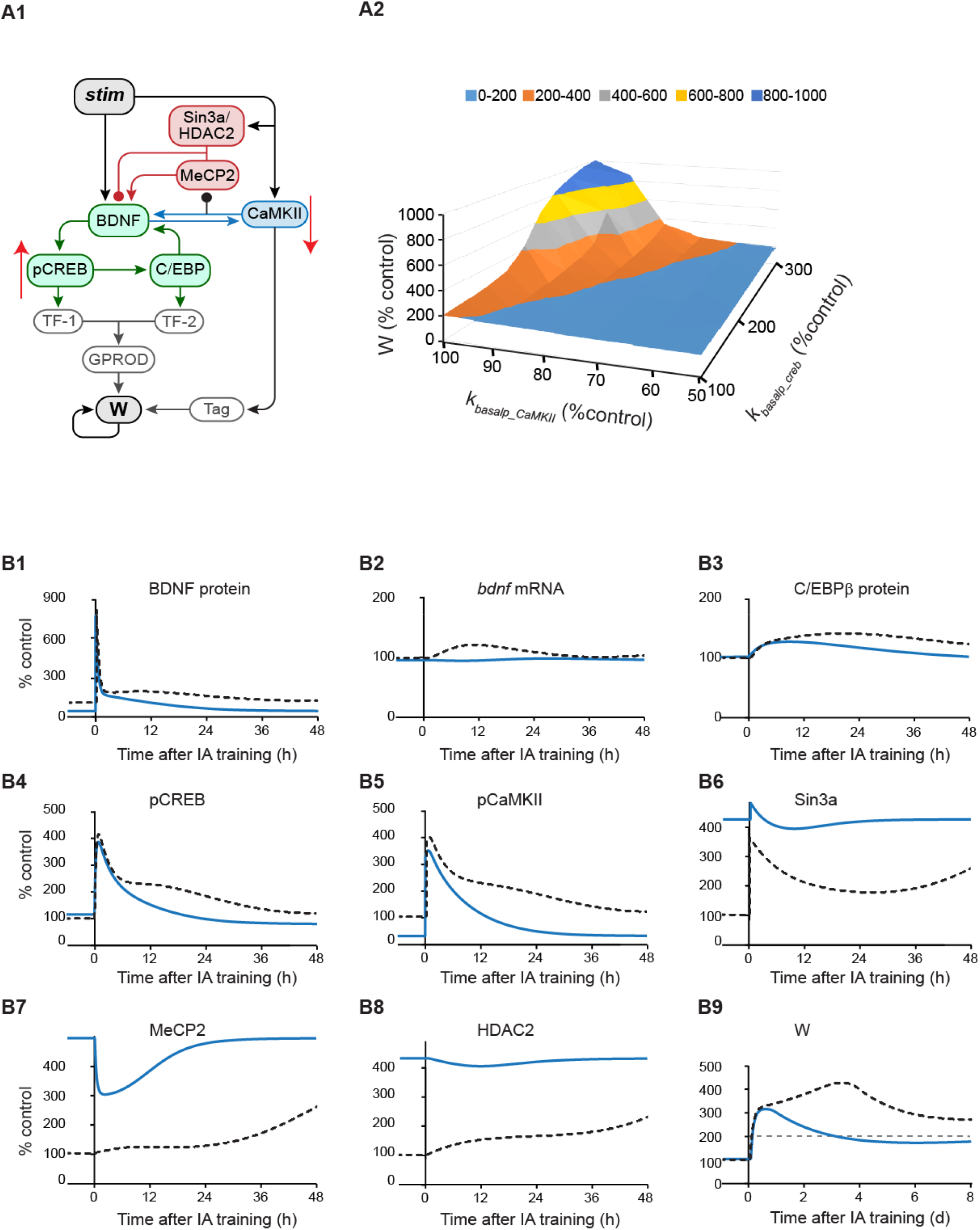
Simulated effects of increased pCREB combined with decreased pCaMKIIα on synaptic weight W. (A1) Increased basal phosphorylation of CREB, *k_basalp_creb_*, concurrent with decreased basal phosphorylation of CaMKIIα, *k_basalp_CaMKII_* (red arrows). (A2) 3D plot of synaptic weight W at day 7 after training with *k_basalp_creb_* increasing from the standard value in Table 1 to 300% of the standard value, and *k_basalp_CaMKII_* decreasing from the standard value in Table 1 to 50% of the standard value, with increased binding of MeCP2, Sin3a and HDAC2 (A2). The light blue area represents W less than 200% of basal level. (B) Example of dynamics of BDNF protein/mRNA (B1-2), C/EBPβ protein (B3), pCREB (B4), pCaMKIIα (B5), Sin3a binding (B6), MeCP2 binding (B7), HDAC2 binding (B8), and W (B9), with the standard parameter values in Table 1 (black dashed) or with *k_basalp_creb_* increased by ~50% and *k_basalp_CaMKII_* decreased by ~50% from control combined with higher binding levels of MeCP2, Sin3a and HDAC2 (blue). Gray dashed line in (B9) represents W at 200% control.

Figure 2B illustrates an example of a simulation when the basal phosphorylation rate of CaMKIIα was decreased by ~50% and the basal phosphorylation rate of CREB was increased by ~50%, relative to control values (Table I). Due to the decrease of basal pCaMKIIα prior to stimulus, releasing its inhibitory effect on MeCP2 binding; MeCP2, Sin3a and HDAC2 binding were all increased to higher basal levels compared to the control simulation (Figs. 2B6-8, blue *vs*. black curves). Because of the higher level of bound MeCP2/Sin3a/HDAC2 complex, BDNF protein and *bdnf* mRNA remained lower than in the control simulation using the standard values of *k_basalp_creb_* and *k_basalp_CaMKII_* in Table 1 (Figs. 2B1-2, blue *vs*. black-dashed curves), and the BDNF-C/EBPβ feedback loop was not fully activated after stimulus (Figs. 2B1-4, blue *vs*. black-dashed curves). W only transiently increased, passing the threshold of 200%, but decreasing to below threshold at day 7 (Fig. 2B9, blue *vs*. black-dashed curves). Taken together, these simulations suggest that enhanced binding of a MeCP2, Sin3a and HDAC2 complex to the *bdnf exon IV* promoter, caused by decreased pCaMKIIα in infant animals, might contribute to rapid forgetting of infantile memory. On the other hand, W did not return to the basal level at day 7 (Fig. 2B9), which suggests that the memory trace is not completely lost, leaving the possibility of memory reinstatement at later times by specific protocols.

### Simulated dynamics of resistance of W to protein synthesis inhibition (PSI)

Injection of the PSI anisomycin into rat dorsal hippocampus blocks >80% of protein synthesis for up to 6 h [5,36]. Thus, we reduced the synthesis rates of BDNF, C/EBP, TF1, TF2, GPROD and W by 80% for 6 h to simulate PSI (Fig. 3A1, red Xs). To investigate the dynamics of memory resistance with respect to the time of PSI addition, PSI was initiated from 0 h to 48 h after the stimulus. The curve of resistance of W in response to PSI (Y axis of Fig. 3B) was built by measuring the degree of PSI-induced reduction of W at day 2 or day 7 post-stimulus (Fig. 3B). The Y axis of Fig. 3B varies from 0% to 100% of the control W in the absence of PSI. If W was reduced to zero, then the reduction was 100%. If W remained intact, then the reduction was 0%. If the attenuation of W in response to PSI decreased, the resistance of W to PSI increased. As expected, early application of PSI was most effective in attenuating W, and the resistance of W to PSI increased with the time of PSI application, leading to an overall decrease in the reduction of W by PSI with time (Fig. 3B). However, unexpectedly, the PSI-induced attenuation at day 2 and day 7 did not show a continuous decrease over time, instead displaying multiple phases. During the first phase after stimulus, when PSI was initiated between immediately and 25 h post stimulus, the reduction of W at day 2 and 7 decreased with respect to the time at which PSI was initiated (arrows #1, Fig. 3B), corresponding to an increase of resistance. During the second phase, when PSI was initiated between 25 and 35 h post stimulus (arrows #2, Fig. 3B), attenuation of W at day 2 remained around the same level (blue curve, Fig. 3B), whereas attenuation of W at day 7 increased with respect to the time at which PSI was initiated (red curve, Fig. 3B), corresponding to a decrease of resistance. After 35 h post-stimulus, attenuation of W at days 2 and 7 resumed decreasing with respect to the time at which PSI was initiated (arrows #3, Fig. 3B). Thus, there is a period of time, between 25 and 35 h post-stimulus, during which the resistance of late W (at 7 days) paradoxically decreases with respect to the time of PSI application. These dynamics illustrate distinct periods during which memory consolidation is differentially affected with respect to the time of PSI application.

**Figure 3.**
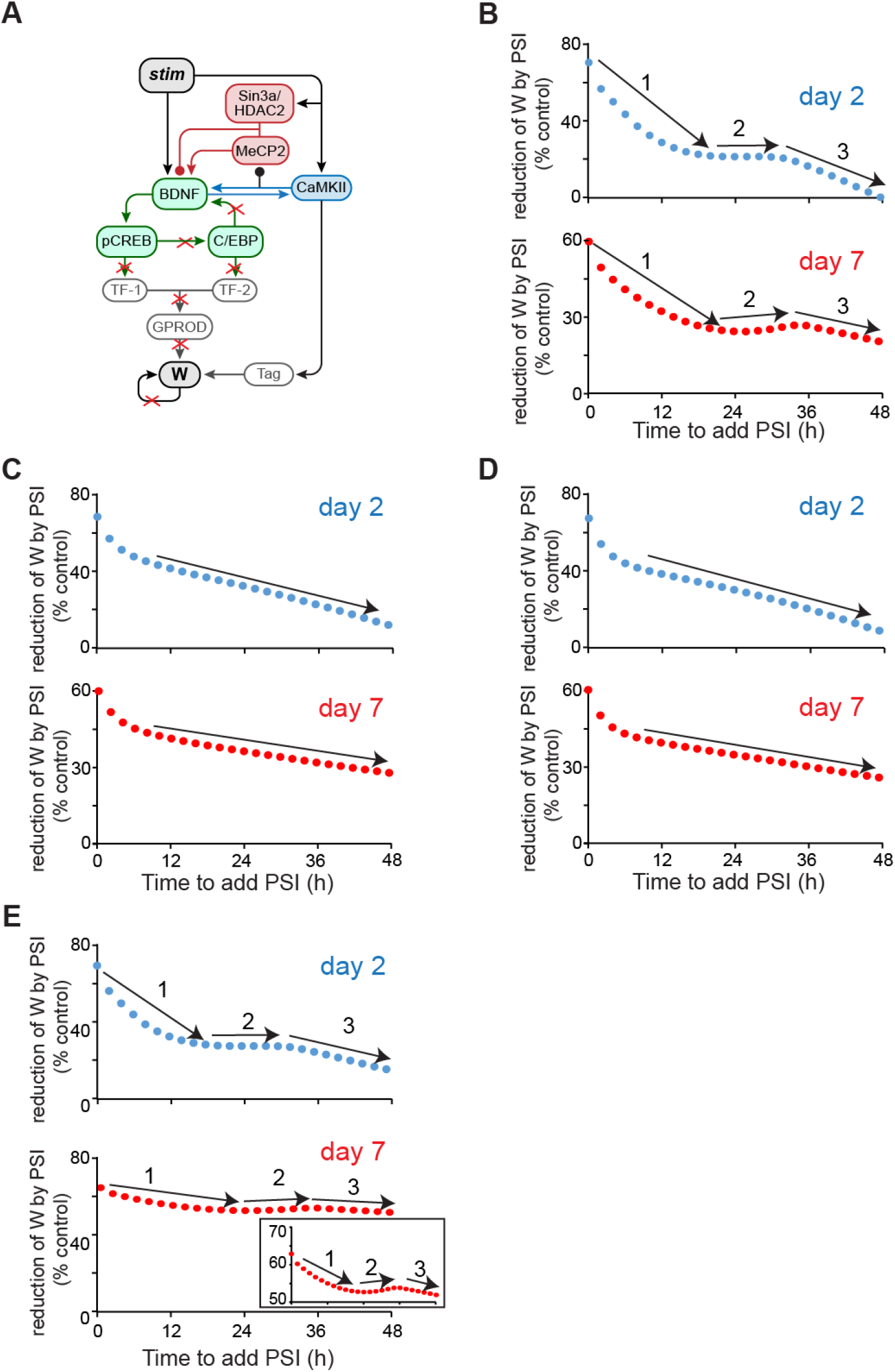
Simulated response to PSI. (A) Schematic model of the pathways that are blocked by PSI (red X’s). (B) Reduction of synaptic weight W at day 2 (blue curve) and 7 (red curve) after training, with the addition of PSI initiated at varying times. ‘1’,’2’,’3’ represent different phases. (C-D) Simulated reduction of W response to PSI in the absence of the BDNF-CaMKIIα feedback loop (C) or of the BDNF-C/EBPβ feedback loop (D) or of the downstream W loop (E) at day 2 (blue) and day 7 (red). Insert in (E), same as main panel except Y axis scale is expanded to more clearly illustrate vertical variations in W.

To test the effects of the BDNF dependent feedback loops on the multiple phases of W reduction in response to PSI, the simulations were repeated, blocking either the BDNF-CaMKIIα feedback loop (Fig. 3C) or the BDNF-C/EBPβ feedback loop (Fig. 3D). Blocking either feedback loop eliminated the second phase during which the reduction of W did not significantly decrease with the time of PSI, for W at day 2 (Figs. 3C-D, blue curves) and day 7 (Figs. 3C-D, red curves). Instead, attenuation of W continuously decreased. Thus, both BDNF dependent feedback loops are necessary for generating multiple phases of W resistance to PSI.

To test the effects of the independent downstream W positive feedback loop on the multiple phases of W resistance in response to PSI, the simulations of Figs. 3B were repeated in Fig. 3E, blocking the downstream W feedback loop. Blocking this loop enhanced the reduction of W by PSI. However, the three phases of resistance of W in response to PSI were still present (arrows #1, 2, 3, Fig. 3E). Thus, this characteristic dynamics of the resistance of W with respect to the time to initiate PSI are not dependent on the downstream W feedback.

### The role of dual effects of MeCP2 on expression of BDNF in the complex dynamics of resistance of W in response to PSI

We compared the time courses of model variables, in the presence *vs*. absence of PSI applied at different times, in order to further examine the mechanism underlying the distinct dynamics of W reduction with respect to the time of PSI initiation during the second phase (Fig. 4). We selected the time points of 25 h and 35 h post stimulus because these two time points are the beginning and end of the second phase (Fig. 3B). Here we investigated why W at day 7 with PSI added at 25 h was greater than W with PSI added at 35 h, (i.e., the reduction of W by PSI applied at 25 h was less than the reduction of W by PSI applied at 35 h). With PSI added 25 or 35 h post stimulus, BDNF decreased in both cases, leading to a decrease in pCaMKIIα (Figs. 4A, 4E, blue *vs*. red *vs*. black dashed curves), releasing its inhibitory effect on MeCP2 binding (Fig. 4G, blue *vs*. red *vs*. black dashed curves). However, when PSI was added at 25 h, MeCP2 binding quickly increased (Fig. 4G, blue arrow), faster than the increase of Sin3a and HDAC2 binding which remained low for several hours before their increase (Figs. 4F, 4H, blue arrows). Thus, MeCP2 alone bound to *bdnf* exon IV promoter to increase the expression of *bdnf*, before Sin3a and HDAC2 could bind it to form the inhibitory complex. The increase in *bdnf* expression occurred after PSI was removed, due to the increased binding of MeCP2 alone (Fig. 4B, blue arrow). Translation of excess mRNA generated a late increase of BDNF that enhanced the recovery of the BDNF-dependent feedback loops, including BDNF, pCREB1 and pCaMKIIα (Figs. 4A, D, E, blue curves), which led to a second wave of increase in Tag and GPROD (Figs. 4I and 4J, blue arrows). The enhanced Tag and GPROD subsequently increased W and its precursor PP (Fig. 4K, blue arrow), with PP initiating the downstream feedback loop, leading to a second wave of increase in W (Fig. 4L, blue arrow). In contrast, when PSI was added later, 35 h post-stimulus, Sin3a and HDAC2 binding were already increasing (Figs. 4F and 4H, red arrows), so that increased MeCP2 binding was directed more to the MeCP2/Sin3a/HDAC2 complex, repressing *bdnf*. Therefore, no enhanced recovery was induced in any variables (Figs. 4A-L, red curves) and the late increase of W was diminished. Thus, due to the dual effects of MeCP2 on the BDNF-dependent feedback loops, W at day 7 had a paradoxically higher value when PSI was added at 25 h than when PSI was added later at 35 h post-stimulus (Fig. 4L, blue *vs*. red curves). These dynamics generate the period within which the resistance of W to PSI decreased with respect to the time of PSI application.

**Figure 4.**
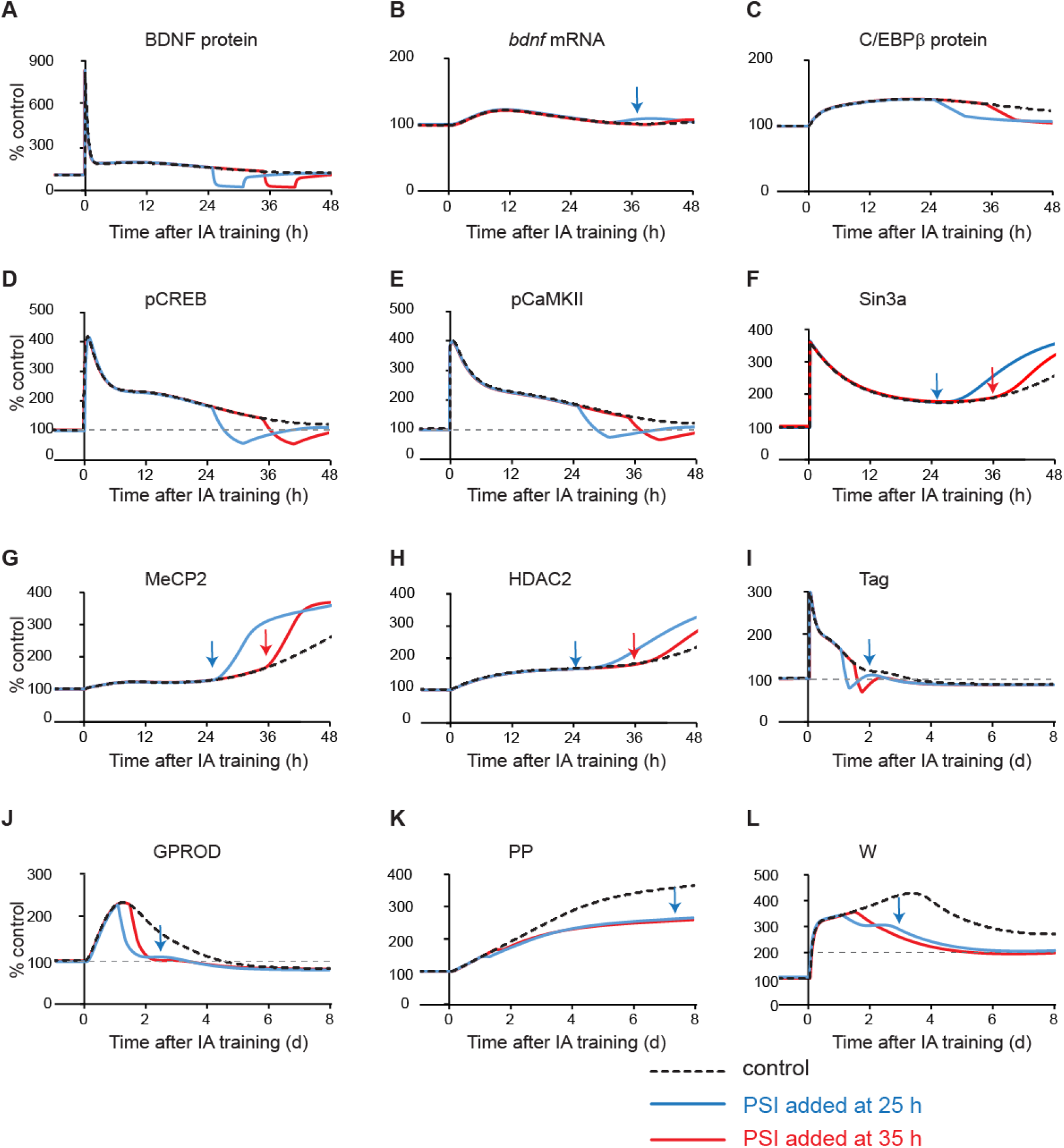
Simulated dynamics of model variables when PSI was added 25 h or 35 h post-stimulus. Example of dynamics of BDNF protein/mRNA (A-B), C/EBPβ protein (C), pCREB (D), pCaMKIIα (E), and Sin3a binding (F), MeCP2 binding (G), HDAC2 binding (H), Tag (I), GROD (J), PP (K), and W (L), without PSI (black dashed) or with PSI added at 25 h post-stimulus (blue), or at 35 h post-stimulus (red).

### Parameter sensitivity analysis to identify other factors that affect the complex dynamics of resistance of W in response to PSI

Parameter sensitivity analysis was performed to examine the extent to which other factors might contribute to the dynamics of resistance of W in response to PSI (Fig. 5). In Fig. 5A, *τ_comp_*, the time constant for the inhibitory effects of the MeCP2/Sin3a/HDAC2 complex on *bdnf* exon promoter, was reduced to 30% of the standard value in Table 1. The second phase of W reduction was eliminated. In Fig. 5B, *τ_E_MeCP2_*, the time constant of activating effects of free MeCP2 on *bdnf exon* promoter, was increased to 300% of standard value in Table 1. The second phase of W reduction was eliminated. Thus, the time scales of the dynamics governing the effects of the MeCP2/Sin3a/HDAC2 complex, and the effects of free MeCP2, played critical roles in the generation of the multiple phases of W reduction in response to PSI. Analogous simulations demonstrated that decreasing *τ_HDAC2_* (the time scale of HDAC2 binding to *bdnf*) to 30% of its standard value or enhancing the response of Sin3a binding to stimulus, *r_sin3a_*, by 300% eliminated the second phase of W reduction in response to PSI (Figs. 5C-D).

**Figure 5.**
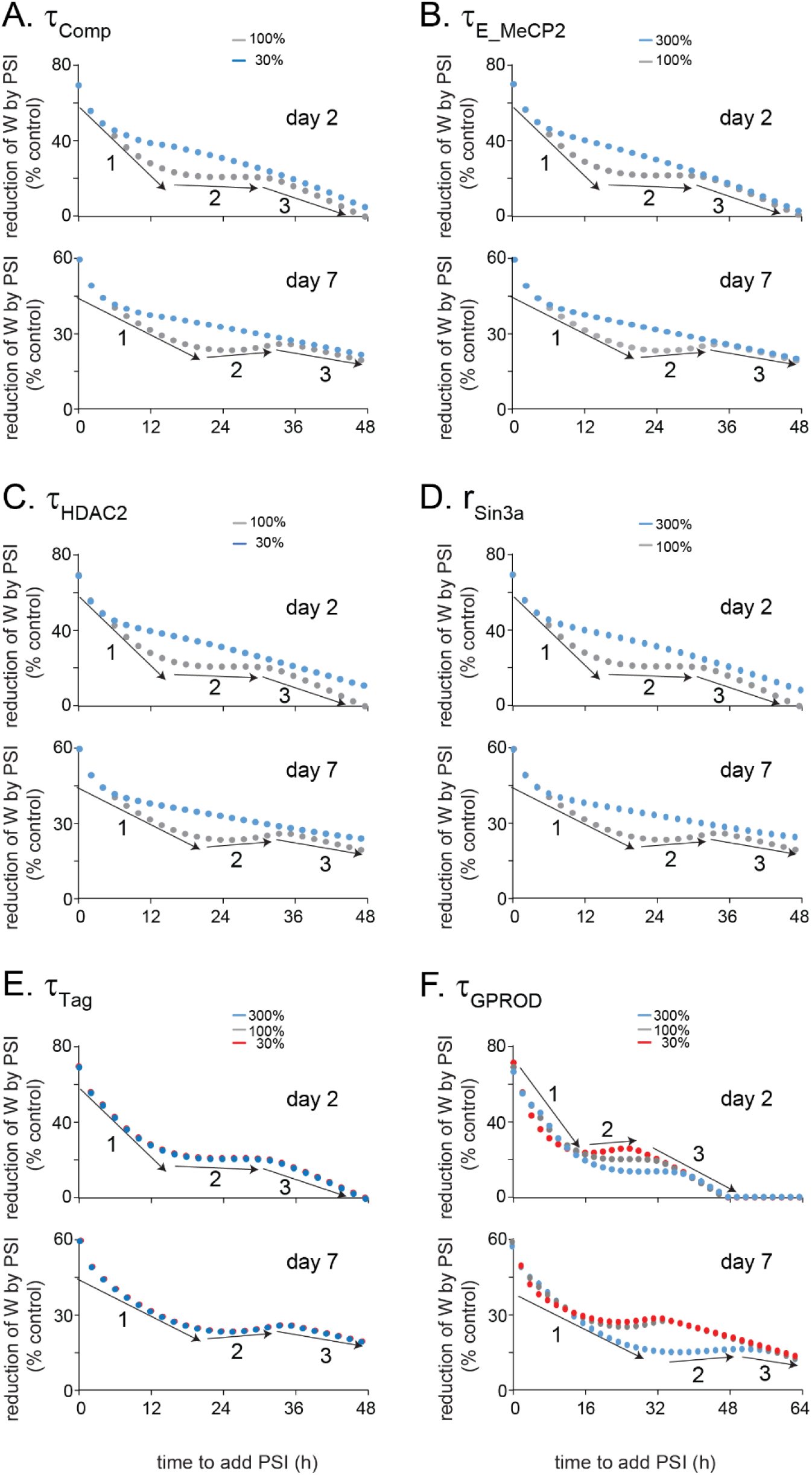
Simulated effects of various parameters on the multiple phases of W resistance in response to PSI. The dynamics of reduction of W at day 2 and 7 post-stimulus, with the addition of PSI initiated from 0 h to 48 h, varied when altering *τ_Comp_* (A), *τ_E_MeCP2_* (B), *τ_HDAC2_* (C), *r_Sin3a_* (D), *τ_Tag_*(E), *τ_GPROD_* (F). 1’,’2’,’3’ represent different phases.

In Fig. 5E, varying *τ_Tag_*, the time constant of Tag activation, from 30% to 300% of the standard value, did not substantially change the multiple phases of W reduction in response to PSI. Likewise, in Fig. 5F, varying *τ_GPROD_*, the time constant of GPROD production, from 30% to 300% of its standard value, also did not substantially change the multiple phases of W reduction in response to PSI. This sensitivity analysis suggests that the variables of the tagging and capture system did not play important roles in the generation of the multiple phases of W reduction in response to PSI. The parameters governing the multiple phases of W reduction are those related to the dynamics of MeCP2, Sin3a and HDAC2 effects on *bdnf* expression.

The simulations also suggest that although BDNF-dependent feedback loops are necessary for the multiple phases of W reduction response to PSI (Figs. 3C-D), they are not sufficient in the absence of further parameter constraints. The multiple phases of response of W only occur within a specific range of the time constants governing the dynamics of regulation of *bdnf* expression by MeCP2, Sin3a and HDAC2. Thus MeCP2, Sin3a and HDAC2 binding kinetics may constitute potential targets for rescuing memory impaired by disruptions.

## Discussion

Empirical studies indicate that recurrent rounds of *de novo* protein synthesis are required to maintain hippocampal memory formation and consolidation for a week or longer [5, 25-26]. However, the ways in which the dynamics of these molecular components of consolidation correlate with different temporal domains of memory is not well understood. BDNF, with levels elevated by feedback, may enhance the expression of downstream proteins such as C/EBP, rendering memory consolidation resistant to disruption during specific post-training periods. In Zhang et al. [3], we developed a differential-equation based model to describe this feedback loop. In the current study, the model of Zhang et al. [3] was revised to further investigate the roles of a BDNF-CREB-C/EBPβ feedback loop and of a BDNF-CaMKIIα dependent feedback loop in the development of resistance to PSI during memory consolidation.

A characteristic feature of a positive feedback loop is that if PSI is not sufficient to reduce the protein synthesis rate to below a threshold for substantial time, then the positive feedback loops will fully recover after PSI is removed [24, 36]. In the model of Zhang et al. [24], after the feedback loops were initiated and the strength grew with time, the resistance of W to PSI was gradually enhanced with time. However, empirical studies suggest that, for some common learning protocols, the resistance of memory to PSI does not continuously increase with the delay in the time to initiate PSI [25-26, 35, 45-47]. In some periods, the resistance decreases with respect to the time of PSI initiation. In this study, we used the revised model to investigate the dynamics of variation of synaptic weight W with respect to the time of PSI. We found multiple phases of W resistance in response to PSI (Fig. 3B). The dual effects of MeCP2 on the expression of *bdnf* helped generate a specific period during which the resistance of W at day 7 post-stimulus to PSI decreased with respect to the time of PSI application.

When PSI was added, the synthesis of BDNF was reduced, leading to a decrease of pCaMKIIα, which subsequently increased binding of MeCP2 to the *bdnf* exon *IV* promoter (Fig. 4). If binding of MeCP2 increased before the increase of Sin3a and HDAC2 binding, MeCP2 alone increased the expression of *bdnf*, enhancing the ability of the BDNF feedback loops to recover after PSI was removed. In contrast, if binding of MeCP2 to the *bdnf* promoter increased together with increased Sin3a and HDAC2 binding, then the MeCP2/Sin3a/HDAC2 complex would inhibit the expression of *bdnf*, suppressing the ability of BDNF feedback loops to recover after PSI was removed. Thus, the multiple phases of W in response to PSI application depend on the temporal difference in the dynamics of binding of MeCP2, Sin3a and HDAC2, *vs.* binding of MeCP2 alone, to regulate the expression of *bdnf* (Figs. 4-5). Simulations also suggest that both BDNF-related feedback loops, but not the downstream W feedback, are required for the multiple phases of W resistance in response to PSI (Fig. 3). The dependence of these phases of response of W to PSI on the dual regulation of *bdnf* by MeCP2 suggests that the special MeCP2 binding kinetics may provide a possible explanation of complex effects of MeCP2 on gene expression [28-29, 48] and a potential therapeutic target for rescuing memory impaired by disruptions.

Dual regulation of *bdnf* by MeCP2 might also help explain apparent rapid forgetting of infantile memory. Multiple hypotheses have been proposed to explain the mechanism of infantile amnesia and reinstatement of latent memory traces [8, 10, 43-44, 49]. In Travaglia et al. [12], in infantile rats, the basal level of pCaMKIIα in dorsal hippocampus is substantially lower than that of adult rats, whereas the basal level of pCREB in dorsal hippocampus is substantially higher than that of adult. We therefore simulated a stimulus response given a decreased basal phosphorylation rate of pCaMKIIα and increased basal phosphorylation rate of pCREB (Fig. 2). Increased pCREB led to an increase of C/EBPβ, tending to augment the BDNF-C/EBPβ feedback loop. However, the decreased pCaMKIIα led to an increased level of the MeCP2/Sin3a/HDAC2 complex, suppressing the BDNF-C/EBPβ feedback loop. The net result of these effects was that although the synaptic weight W post-stimulus transiently increased due to the increase in pCREB, this increase was not sufficient to overcome the suppression of feedback and maintain W at day 7 post-stimulus at a sufficiently high level to retrieve memory (Fig. 2B, blue curves). However, W did remain at an elevated level, compared to pre-stimulus baseline, at day 7, suggesting memory reinstatement due to specific protocols may be elicited at later times. The simulation results also help explain why exogenous application of BDNF can restore infantile memory [9]. It is possibly via overcoming the suppression of *bdnf* expression by MeCP2/Sin3a/HDAC2 complex. Multiple signaling cascades in addition to those modeled here are involved in infantile amnesia. Nevertheless, these simulations suggest that in infant animals, decreased basal CaMKIIα activity and increased basal levels of bound MeCP2/Sin3a/HDAC2 complex may contribute to fast forgetting of infantile memory in the early phase.

Although the revised model is based on some relatively speculative assumptions, several predictions can be derived from the present simulations. First, the simulations predict critical roles of MeCP2 in helping to delineate the multiple phases of resistance of W to PSI. Second, the model also predicts ways in which the dynamics of W and other variables would be altered by changing kinetics of the BDNF-related feedback loops, or binding of the transcription factors MeCP2, Sin3a, or HDAC2 to the *bdnf exon* promoter. Finally, the model also predicts that decreased activation of CaMKIIα, combined with increased MeCP2/Sin3a/HDAC2 binding, underlies in part the rapid forgetting of infantile memory. If these predictions can be validated by empirical studies, they may suggest paths towards novel potential therapeutic targets for rescuing memory impairment.

